# Extensive *de novo* mutation rate variation between individuals and across the genome of *Chlamydomonas reinhardtii*

**DOI:** 10.1101/015693

**Authors:** Rob W. Ness, Andrew D. Morgan, Radhakrishnan B. Vasanthakrishnan, Nick Colegrave, Peter D. Keightley

## Abstract

Describing the process of spontaneous mutation is fundamental for understanding the genetic basis of disease, the threat posed by declining population size in conservation biology, and in much evolutionary biology. However, directly studying spontaneous mutation is difficult because of the rarity of de novo mutations. Mutation accumulation (MA) experiments overcome this by allowing mutations to build up over many generations in the near absence of natural selection. In this study, we sequenced the genomes of 85 MA lines derived from six genetically diverse wild strains of the green alga *Chlamydomonas reinhardtii*. We identified 6,843 spontaneous mutations, more than any other study of spontaneous mutation. We observed seven-fold variation in the mutation rate among strains and that mutator genotypes arose, increasing the mutation rate dramatically in some replicates. We also found evidence for fine-scale heterogeneity in the mutation rate, driven largely by the sequence flanking mutated sites, and by clusters of multiple mutations at closely linked sites. There was little evidence, however, for mutation rate heterogeneity between chromosomes or over large genomic regions of 200Kbp. Using logistic regression, we generated a predictive model of the mutability of sites based on their genomic properties, including local GC content, gene expression level and local sequence context. Our model accurately predicted the average mutation rate and natural levels of genetic diversity of sites across the genome. Notably, trinucleotides vary 17-fold in rate between the most mutable and least mutable sites. Our results uncover a rich heterogeneity in the process of spontaneous mutation both among individuals and across the genome.

## Introduction

Understanding the processes that generate new genetic variation from mutation is a key goal of genetics research. It is widely believed that the majority of new mutations that affect functional elements of the genome are deleterious. In humans, new mutations cause Mendelian genetic disorders, play a direct role in polygenic disease (e.g. Veltman and Brunner 2012), and are a major factor in cancers (e.g. Alexandrov et al. 2013a). New mutations also play a central role in evolutionary biology, since the variation that fuels adaptive evolution is ultimately derived from advantageous mutations. For example, the input of new variation from mutation is pivotal for theory to explain the evolution of recombination and sex (reviewed in Otto 2009).

If new mutations are harmful, theory predicts that the mutation rate should evolve towards zero, because individuals with higher mutations rates will suffer a greater mutational load. However, the mutation rate is always greater than zero in nature, ranging over seven orders of magnitude (reviewed by Drake 2006), and two main explanations have been proposed for this. One explanation is that there is a limit to the fidelity of DNA repair, due to a trade-off between the benefit of further reducing the mutation rate and the costs of increased fidelity (Kimura 1967). Alternatively, a ‘selection-drift’ barrier may constrain progress toward lower mutation rate when the selective advantage of further improvement becomes so small that new mutations decreasing the mutation rate are effectively neutral (Lynch 2010). Evidence for a selection-drift barrier comes from the negative correlation between the mutation rate per generation and effective population size (N_e_) (Sung et al. 2012). However, when mutation rate is expressed per cell division, there is much less variation between species and little relationship with N_e_, consistent with the constraint on the fidelity of replication hypothesis. It is currently difficult to fully evaluate the support for these hypotheses, however, because studies of mutation are restricted to a small number of taxa, few genotypes per species and a limited number of mutation events.

Although there is clear evidence for variation between species, we know relatively little about the extent of mutation rate variation within species. Individuals with unusually high mutation rate have been isolated from natural populations of prokaryotes (Matic et al. 1997; Sundin and Weigand 2007), but no natural mutators have been found in eukaryotes. This discrepancy likely stems from the fact that prokaryotes are asexual whereas eukaryotes are predominantly sexual. Theory predicts that in an asexual population, a mutator allele can hitchhike to high frequency if it causes a beneficial allele on the same genetic background (Johnson 1999). In contrast, recombination in sexual populations uncouples a mutator from a linked beneficial allele, so the mutator allele is then expected to be selected against because of its association with linked deleterious mutations (reviewed by Drake et al. 1998). Although a smaller amount of mutation rate variation is expected in sexual than asexual species, mutations that alter the mutation rate are nevertheless expected to occur, and potentially provide the basis for mutation rate evolution. Mutation rate variation within a species may also reflect mutation-selection balance, whereby new deleterious alleles that alter the mutation rate continually arise and are purged by selection. In this scenario, intraspecific mutation rate variation will reflect the distribution of phenotypic effects of mutations that alter DNA repair and stability and the effectiveness of selection against them. In the largest study of spontaneous mutation in humans, there was little evidence for mutation rate variation among individuals after accounting for parental age (Kong et al. 2012). Father’s age was also an important factor explaining mutation rate variation in chimpanzees (Venn et al. 2014). Similarly, there was no evidence of mutation rate variation between two strains in both *Caenorhabditis elegans* and *C. briggsae* (Denver et al. 2012). There is evidence from *Drosophila* that individuals in poor condition have elevated mutation rates (Sharp and Agrawal 2012) and a separate study comparing two inbred lines revealed a 2.4-fold difference in the rate of mutation (Schrider et al. 2013). Moreover, two independent experiments in *Chlamydomonas reinhardtii* suggested that there is a 5-fold difference in the mutation rate between two natural strains (Ness et al. 2012; Sung et al. 2012).

In addition to mutation rate variation within and between species, there is also evidence that mutation rate varies across the genome. Such heterogeneity is expected to alter the rate of evolution across the genome and to create variation in the susceptibility of genes or sites to deleterious or beneficial mutations. There is clear evidence for fine-scale variation in the rate of mutation. At the scale of individual sites, G:C positions tend to mutate at higher rates than A:T positions, and transitions from G:C→A:T are the most common change in a broad range of species (for example bacteria (Hershberg and Petrov 2010), animals (Kong et al. 2012; Schrider et al. 2013), fungi (Zhu et al. 2014) and plants (Ness et al. 2012)). Similarly, the bases surrounding a mutated site have a strong effect on mutability. For example, the high frequency of G:C→A:T transitions in mammals is driven by the deamination of methylated C_p_G sites (Ehrlich and Wang 1981). In general, the bases flanking a particular site, referred to as the ‘sequence context’, are one of the best predictors of mutation rate (Michaelson et al. 2012; Neale et al. 2012; Samocha et al. 2014; Zhu et al. 2014). However, the underlying mechanisms and the consistency of the effect of sequence context on mutability across species is unknown.

At a broader genomic scale, evidence for mutation rate heterogeneity is weaker. Sequencing of MA lines in S. *cerevisiae* (Zhu et al. 2014) and *D. melanogaster* (Schrider et al. 2013) found no evidence of mutation rate variation between chromosomes. Although there is evidence that mutation rate increases as a function of replication timing (Stamatoyannopoulos et al. 2009; Lang and Murray 2011), this finding has not been supported by direct estimates of mutation rate (Samocha et al. 2014; Zhu et al. 2014). A variety of other genomic properties have been linked to increased susceptibility to mutation, including transcription level, nucleosome occupancy, DNAse hypersensitivity and recombination rate (e.g. Michaelson et al. 2012). If these factors strongly influence mutation and generate variation between sites or large scale patterns of mutation rate variation, it is important to quantify their effects, in order to facilitate better predictive models of DNA sequence evolution.

Detailed investigations of the process of spontaneous mutation and the extent of mutation rate variation are limited. This is because spontaneous mutations are very rare, constraining direct observation of sufficient numbers of mutations to infer the underlying biology. Sequencing of parents and their offspring is an increasingly common method for directly identifying de novo mutations (e.g. Keightley et al. 2014a; Keightley et al. 2014b). Although this approach has advantages, it is currently very expensive to sequence enough families to observe large numbers of mutations and has therefore only been applied on a large scale in humans (Kong et al. 2012). Another approach is to maintain experimental populations for many generations under minimal natural selection to allow mutations to accumulate regardless of their fitness consequences. Increasing the strength of genetic drift by bottlenecking the population each generation allows random, unbiased accumulation of all but the strongest deleterious mutations. These ‘mutation accumulation’ (MA) experiments have been used in a variety of species to investigate the phenotypic effects of new mutations (reviewed in Halligan and Keightley 2009) and are now being paired with whole genome sequencing to identify individual mutations. MA studies have generally been limited to sequencing a small number of genomes, and only two studies (Schrider et al. 2013; Zhu et al. 2014) have tested for heterogeneity in mutation rate across the genome, and no study has included more than two ancestral genotypes from a single species. In this study, we sequenced the genomes of 85 MA lines derived from six genetically diverse wild strains of the model green alga *C. reinhardtii.* We identified 6,843 mutations, seven-fold more than any previous MA study, and integrate this data with detailed annotation of genomic properties to investigate the process of spontaneous mutation with unprecedented detail. Specifically, we address the following fundamental questions (1) What is the relative frequency of different kinds of mutation, including the base spectrum and rate of insertion and deletion mutations? (2) What is the extent of mutation rate variation between individuals within a species? (3) Is there evidence of mutation rate heterogeneity across the genome and what genomic properties predict the rate of mutation at individual sites?

## Results

We conducted a mutation accumulation experiment in six genetically diverse wild strains of *C. reinhardtii* that were chosen to broadly cover the geographic range of known *C. reinhardtii* samples in North America (Table 1). 15 replicate MA lines from each of the six ancestral strains were initiated for a total of 90 MA lines. 85 of the initial 90 MA lines survived to the end of the experiment. The mean number of generations per MA line was 940 (range 403 to 1,130). DNA was extracted from each line and sequenced using Illumina whole genome sequencing allowing us to identify mutations in an average of 75.4Mbp per line (72.5% of genome, range 58.5-84.9Mbp; See Materials and Methods for details on mutation calling). In total, we identified 6,843 mutations, including 5,716 single nucleotide mutations (SNMs) and 1,127 short indels. To confirm these mutation calls, we Sanger sequenced a random sample of 192 mutations. 138 were successfully amplified and sequenced, 115 of 117 SNMs were confirmed and 19 of 21 indels were confirmed, resulting in an accuracy of 98.3% and 90.5% for SNMs and indels respectively.

**Table 1.** Ancestral strains of Chlamydomonas *reinhardtii* used for mutation 1 accumulation (MA). Each of the six strains was used to generate 11-15 replicate MA lines. The original sampling location, date and mating type (+/-) are indicated. The total number of single nucleotide mutations (SNMs) and short indels (<50bp) identified across all replicates of each strain are reported, along with the mean number of high quality (‘callable’) genomic sites sequenced in each strain.

| Ancestral Strain | Collection Location/Year | Mating Type | MA lines | Mutations (SNMs / short indels) | Mean callable sites (Mbp) |
| --- | --- | --- | --- | --- | --- |
| CC-1373 | Massachusetts/1945 | + | 12 | 1696/222 | 78.8 |
| CC-1952 | Minnesota / 1986 | - | 14 | 366/66 | 74.4 |
| CC-2342 | Pennsylvania / 1989 | - | 11 | 824/73 | 72.0 |
| CC-2344 | Pennsylvania / 1989 | + | 15 | 946/181 | 75.3 |
| CC-2931 | North Carolina / 1991 | - | 14 | 1215/405 | 72.5 |
| CC-2937 | Quebec / 1993 | + | 15 | 508/149 | 78.6 |

### Mutation rate variation among genotypes

Including all MA lines, the total mutation rate was, μ = 1.15×10^−9^ muts/site×generation, with SNM and indel mutation rates of μ_SNM_ = 9.63×10^−10^ and μ_INDEL_ = 1.90×10^−10^, respectively. The mutation rate varied substantially among the MA replicates and between ancestral strains. Mutation rates of the individual MA lines ranged over nearly two orders of magnitude from μ_CC-1952-MA4_ = 5.65×10^−11^ to μ_CC-2344-MA1_ = 4.94×10^−9^. There was significant variation in the mean mutation rate among the strains (*F_15_* =30.96, P<0.0001, see Fig. 1). Post hoc Tukey tests showed strain CC-1373 had an average mutation rate significantly higher than all other strains (μ=28.1 ×10^−10^, *P 0.01 to <0.001).* This rate was nearly 7-fold higher than CC-1952 (μ=4.05×10^−10^), which had the lowest mutation rate, and was significantly lower than CC-1373 ( P<0.001), CC-2931 (μ=15.6×10^−10^, P<0.001) and CC-2342 (μ=11.1 ×10^−10^, P<0.01). Within strains CC-2344 and CC-2931, there were individual MA lines with significantly higher mutation rates, 3.5×and 8.0×above their respective strain means (μ estimates are outside the 99.99% CI of their ancestral strain mutation rates, μ_cc-2344-MA1_=56.9×10^−10^, CC-2344 CI = 2.6 −12.0×10-^10^; μ_cc-2931-MA5_=36.2×10^−10^, CC-2931 CI = 7.2-20.0×10^−10^).

**Figure 1.**
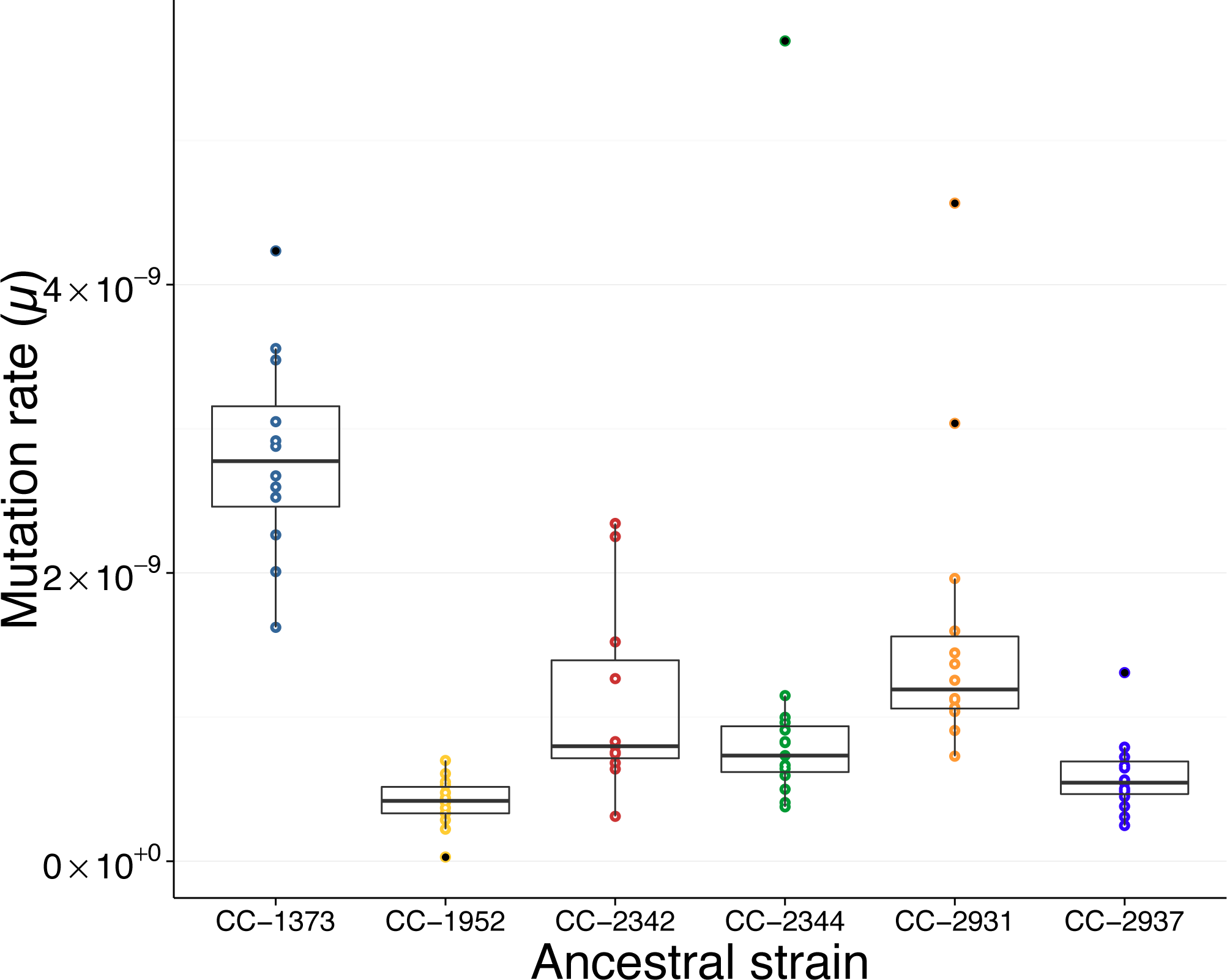
Variation in mutation rate between strains. Total mutation rate (*μ* = total mutations / (site χ generation)) for each of the MA lines, categorized based on their ancestral strain. The boxes outline the 1st to 3rd quartile of the mutation rate in lines from a given ancestral strain, the thick horizontal line indicates the median mutation rate and the whiskers extend to the last data point that is within 1.5× the interquartile range, points outside the whiskers are filled black.

### Indel mutations

There were significantly more short deletions (613) than insertions (514) (χ^2^=8.7, P < 0.005) and these tended to be larger (mean length −7.9 and +5.9, respectively, Mann-Whitney U test, W=112604.5, *P<* 2.2×10^−16^), but the difference was not significant. MA lines of strain CC-2931 had an unusually high number of indels (408) due to an abundance of 9bp deletions. 120 of 408 indels in CC-2931 were 9bp deletions compared to a mean of five 9bp deletions in each of the other strains. These deletions did not appear to have any shared sequence motif nor were they associated with coding exons, repetitive sequence or any genomic property that we could identify. After adjusting for the excess of 9bp deletions in CC-2931 by substituting the mean number of 9bp deletions found in the other strains, there were similar numbers of insertions and deletions, but deletions were still significantly longer (W=100759.5, *P=* 3.3×10^−9^).

### Spatial heterogeneity

When mutation rate was measured in 200kbp sliding windows, μ ranged from 0.0 to 23.5×10^−10^ By comparing the distribution of mutation rates for each window with a simulation distribution, much of this variation could be accounted for as noise around the genome average mutation rate (KS test *D* = 0.038, *P<* = 0.43). In 1,000 simulations where mutation positions were randomized, the 95% confidence interval (CI) was μ = 5.3 - 18.3×10^−10^ compared to a 95% CI of μ = 4.8- 19.4×10^−10^ in the observed data. 8% of 200kbp windows were above the 95th percentile of simulated mutation rates, suggesting a very slight excess of windows with a high mutation rate. Notably, the chloroplast genome had a mutation rate of μ_cpDNA_ = 5.17×10^−9^, nearly 4.5× the genome average.

We detected a significant deviation in the distribution of minimum intermutation distance compared to those expected under simulation (Fig. 2, K-S test: *D* =0.048, *P<* = 4.5×10^−14^). There was a large excess of mutations clustered very near to one another (<100bp apart) and most of this excess was caused by mutations at adjacent sites. Specifically, we expected zero adjacent mutations, but identified 55 mutations where two adjacent sites were mutated, each of which was visually inspected in the Integrated Genomics Viewer, IGV (Thorvaldsdôttir et al. 2012). 27 of these clustered mutations occurred at CC (or GG) sites, and 25 of 27 mutated to AA/AT/TA/TT. We also found a number of indels where a short amount of sequence was replaced by an unrelated stretch of sequence. These complex indels are often reported by GATK HaplotypeCaller as a separate deletion and insertion rather than a single event. The excessive clustering occurred only within MA lines and when we limited our analysis to test for the presence of clustering of mutations found in different lines there was no evidence for this effect (K-S test *D* = 0.02, *P<* 0.13).

**Figure 2.**
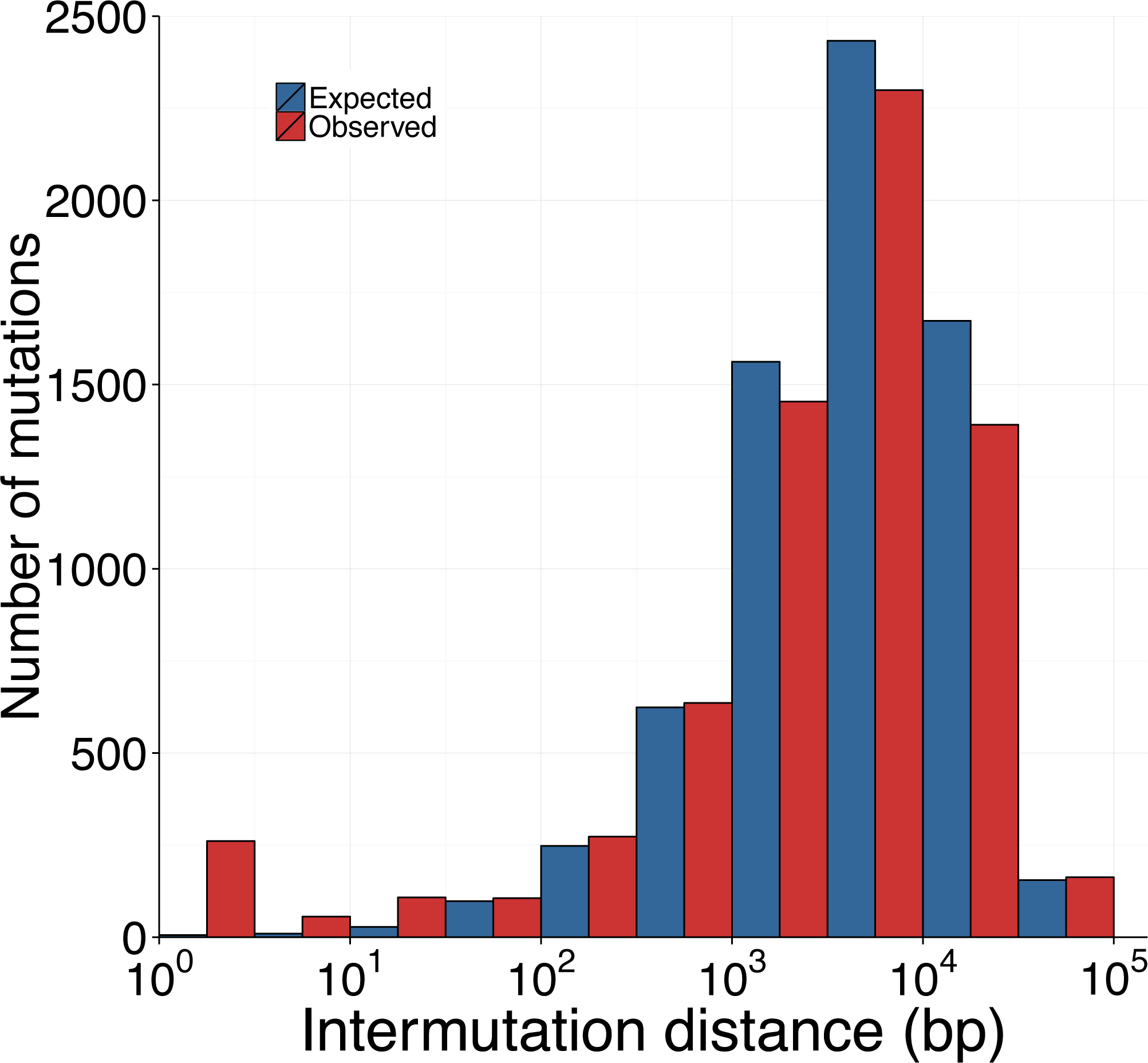
Expected and observed distributions of intermutation distance. Comparison of observed (red) and expected (blue) distributions of the distance between mutations. In this plot, intermutation distance was measured as the nearest mutation irrespective of the MA line or strain it occurred in. The expected distribution was generated by randomizing the location of mutations in each MA line and recalculating the intermutation distances. The simulation was repeated 1000 times and the average of those iterations is shown here.

### Base composition

Treating the strand symmetrically we found a significantly non-random distribution of the six possible SNMs (χ^2^=1630.3, *P* < 0.0001; Fig. 3). Mutations occurring at C:G sites were 4.2×more frequent than mutations at A:T sites, after correcting for genomic base composition, and this pattern was consistent across all MA lines and ancestral strains. Transitions from C:G→T:A were over-represented nearly two-fold compared to random expectation. While transitions from A:T→G:C were more common than the other mutations possible at A:T sites, they were still less common than any mutation at C:G sites. Transversions from A:T→C:G or T:A were the least common and were found 2.4× less frequently than expected by chance.

**Figure 3.**
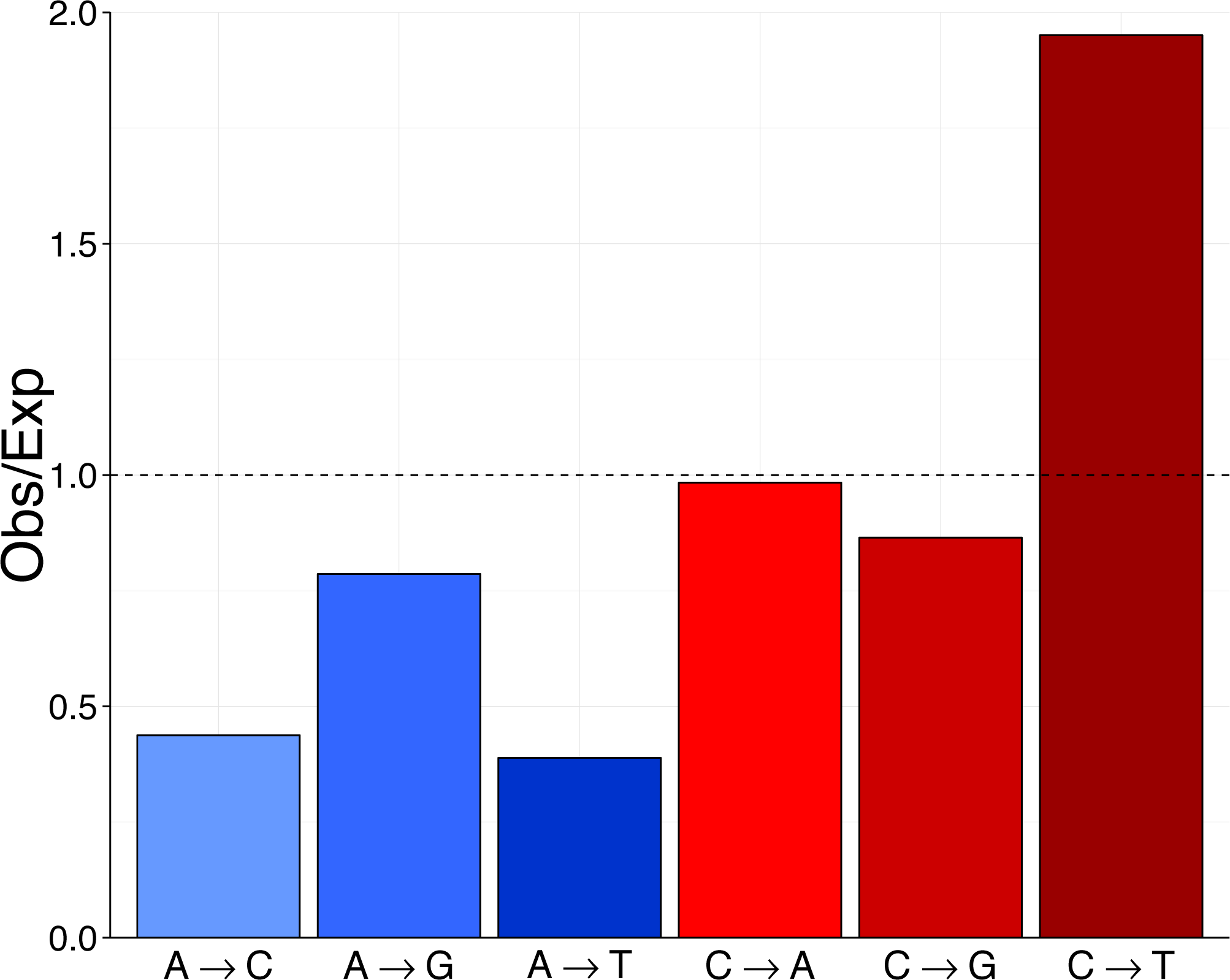
Mutation base spectrum of single nucleotide mutations. Base mutation spectrum of 5716 single nucleotide mutations (SNMs). The deviation of the mutation rate for each of the six possible SNMs relative to its expectation based on equal mutation rates was calculated as the observed number of mutations of each kind divided the number of mutations expected if mutations occurred randomly with respect to base. Background base composition was calculated only from sites that have high quality genotype calls (callable sites).

To assess the effect of the local sequence context on mutation rate, we measured the frequency of the bases surrounding random A:T and C:G sites in the genome and compared this to the base frequencies in the window surrounding SNMs (Fig. 4). We found non-random patterns surrounding all six kinds of mutation, but the extent of the deviation was strongest for mutations at C:G sites. The deviation was particularly strong in the 2-4bp upstream of mutations at C:G sites and to a lesser extent 1bp downstream of all mutation types. Specifically, the composition of the two nucleotides immediately upstream of mutated C:G sites was strongly biased. In the case of the CTC trinucleotide, for example, where the final C was mutated, that mutation rate was 4.5x the background rate.

**Figure 4.**
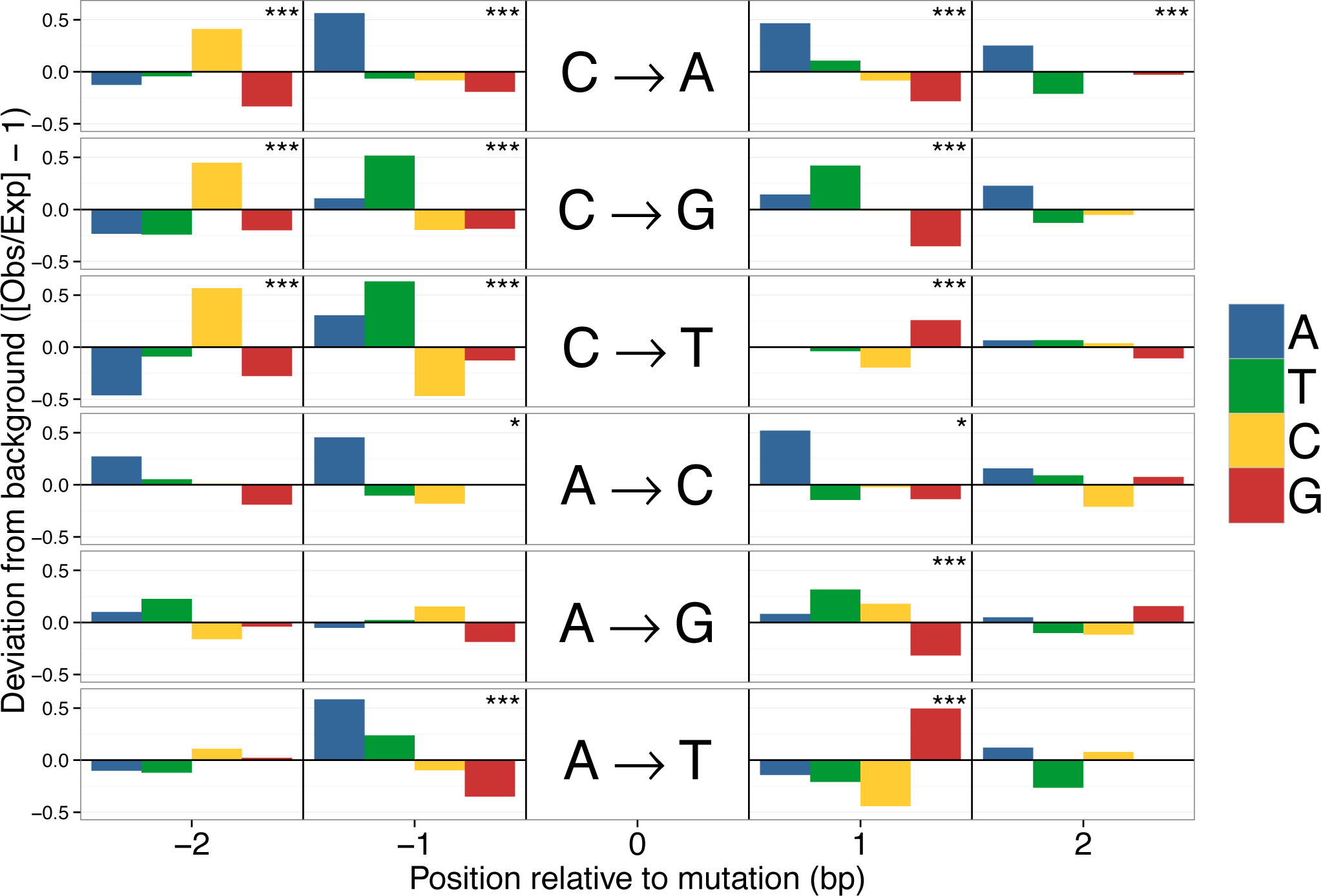
Sequence context of spontaneous mutations. Deviations in the local sequence context of the 2bp flanking mutated sites. Deviations were calculated from the observed frequency of each base (A, T, C, G) in the flanks of mutated sites and the expected background composition based on flanking sequences of 10^6^ random A:T or C:G sites. Each horizontal panel represents one of the six possible mutations indicated in the centre. Significant deviations from the background base composition at each position were detected with tests and indicated as *P <0.05, **P<0.01, *** P<0.001 (alpha-values were adjusted for multiple tests using a Bonferroni correction).

### Mutability

To determine which genomic properties influenced the mutability of individual sites we used logistic regression to differentiate between the identified mutations and randomly selected not mutated sites. Using this model, we then calculated the probability of mutation, or ‘mutability’, for each site in the genome (See Materials and Methods for details). To assess the accuracy of the model we binned sites in the genome based on their mutability (0-1) and calculated the observed mutation rate in each bin (bin width = 0.01). The predicted mutability of sites was strongly correlated with observed mutation rate (Fig. 5, R^2^=0.953, weighted by number of site-generations per bin). To ensure that the fit was not due to using the same mutations to generate the model and assess its fit, we also trained a model using a random subset of 1,000 mutations and excluded these sites when assessing the fit. As with the full data set, predicted and observed mutability were highly correlated (R^2^=0.88). The fit was slightly reduced, presumably because using fewer mutations to calculate mutation rates led to more noise. Although mutability ranged from nearly 0 to 1.0, we found that 99.9% of the genome had mutability values between 0.01 and 0.30, corresponding to mutation rates of 0.25-55.9×10^−10^. The top 25% of genome by mutability accounts for 57% of all mutations. Mutability was highest for sites in the 3’ and 5’ UTRs (predicted μ = 1.37×10^−9^) and lowest for 0-fold and 4-fold degenerate sites (predicted μ= 7.92×10^−10^).

**Figure 5.**
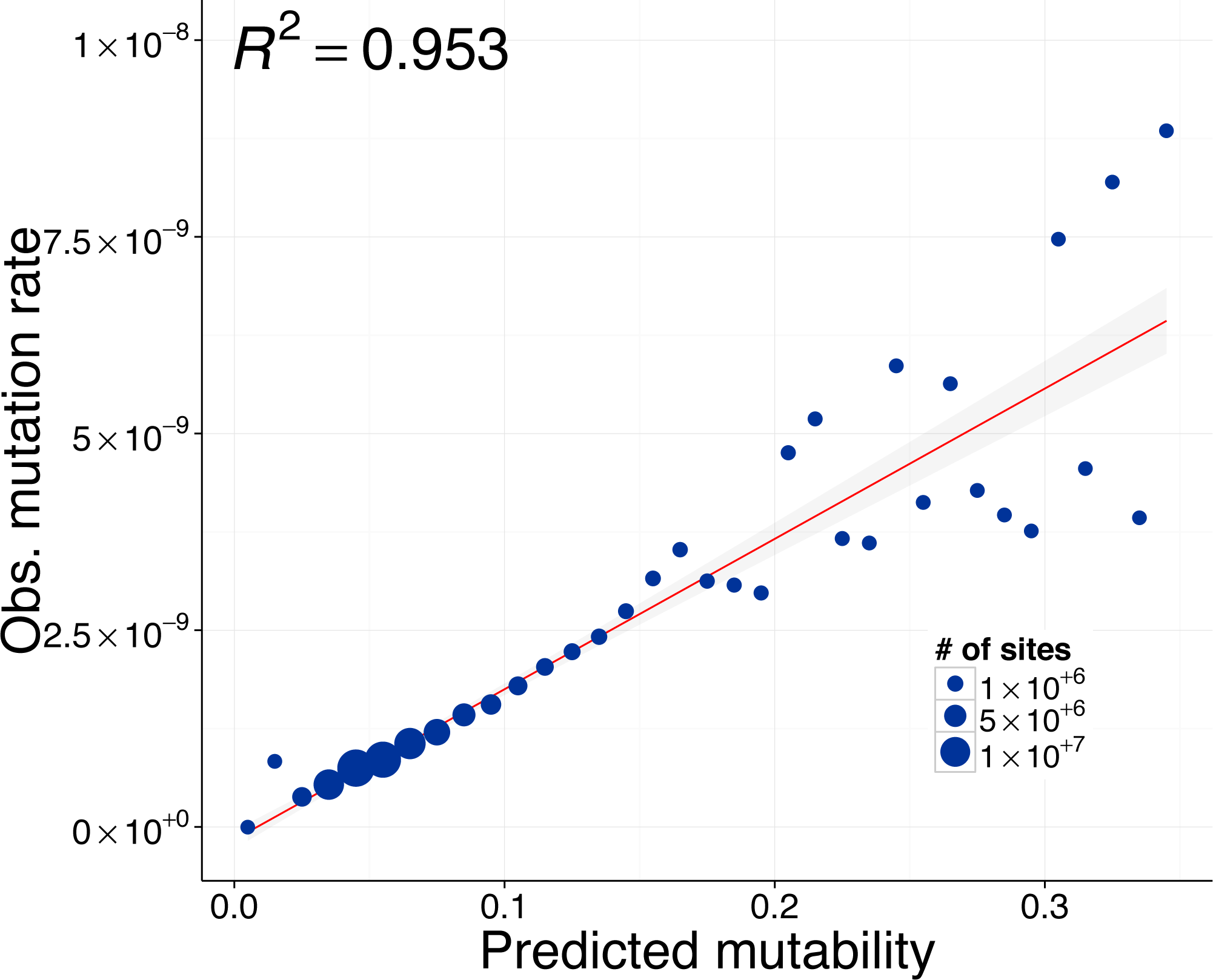
Linear fit between observed mutation rate and predicted mutability. Mutability was estimated using a logistic regression, where the presence or absence of a mutation was the response variable, and a variety of genomic properties were used as predictors (see supplementary table S1). Each point represents multiple genomic sites placed in discrete bins (width = 0.01) based on each site’s mutability score. The size of each point is proportional to the number of sites in the genome with a given mutability. Observed mutation rates for each point were calculated as the number of observed mutations divided by the total number of callable sites-generations in that bin. The linear regression was weighted by the number of sites in each bin and the shaded grey area around the line represents the 95% confidence region.

In neutrally evolving haploid DNA the level of nucleotide diversity (θπ) is expected to be twice the product of mutation rate and the effective population size (2Νβμ), We binned silent sites (intergenic, intronic and 4-fold degenerate sites) into 100 uniformly spaced mutability categories from 0.0-1.0 and calculated θπ for each bin using natural variation in the six ancestral strains used to initiated the MA lines. We found that, as predicted, sites with higher mutability have higher neutral genetic diversity (Fig. 6).

**Figure 6.**
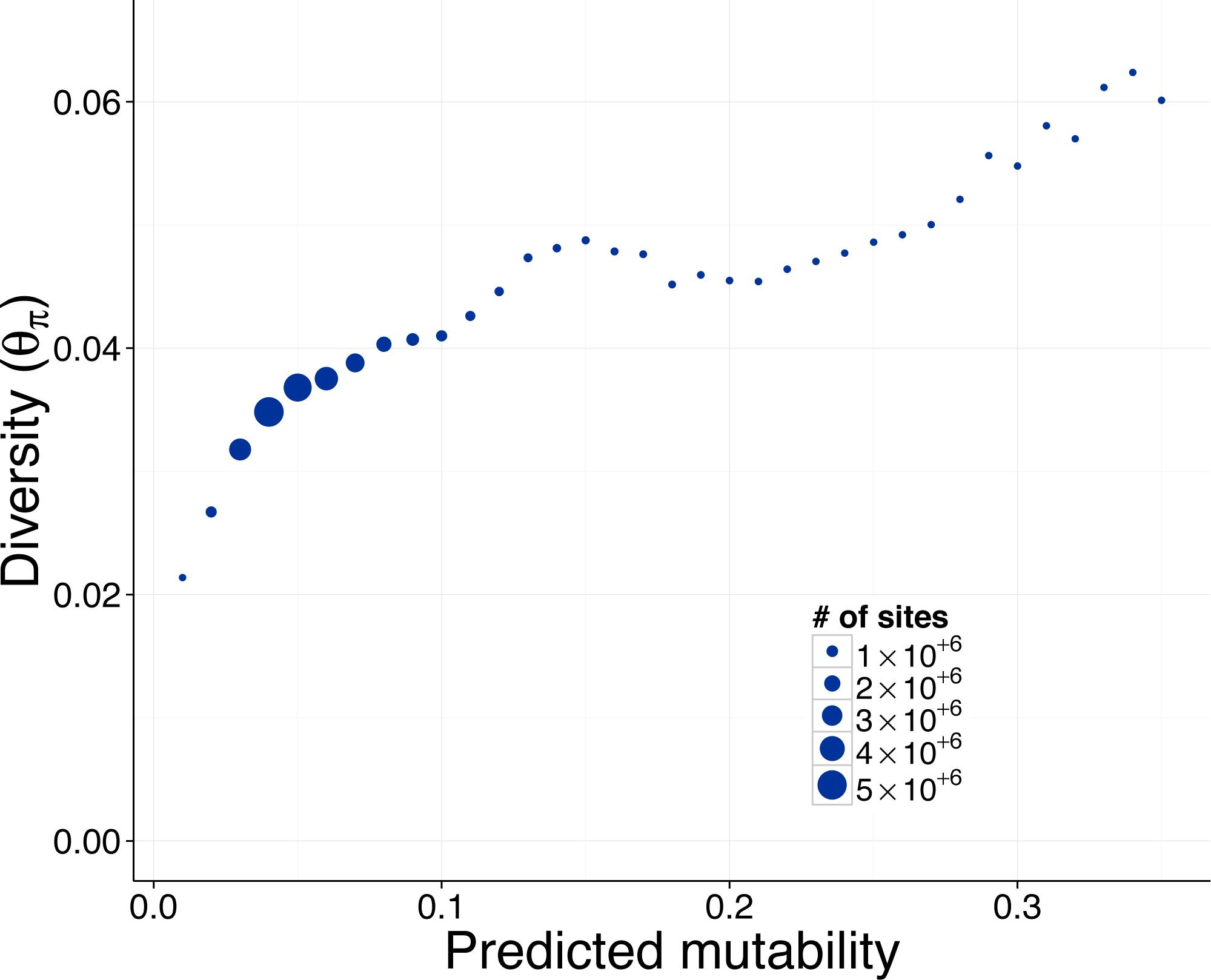
Relationship between natural genetic diversity and predicted mutability. Each point represents multiple genomic sites placed in discrete bins (width = 0.01) based on the predicted mutability of each site. Only putatively neutral sites (intronic, intergenic and 4-fold degenerate sites) were included in this figure. Nucleotide diversity (*θ_π_*) was calculated in each bin from the six ancestral strains used to start the mutation accumulation lines. The size of each point is proportional to the number of sites in the genome with a given mutability.

### Factors influencing mutability

From the model of mutation rate, we extracted the relative contribution of different genomic properties to mutability. To allow comparison among the genomic properties, we scaled continuous predictors so that a change from 0 to 1 was a change of one standard deviation. We found that GC-content of the surrounding genome strongly influenced the mutability at a site. Increasing the GC content of the 10bp surrounding a site increased its mutability (GC% 10bp, odds ratio = 1.38), but at larger scales GC content was negatively related to mutability (GC% 1000bp, odds ratio = 0.12). The negative relationship between GC-content and mutation rate was supported by a highly significant correlation between the observed mutation rate and GC content across the genome (see supplementary Fig. S1, R^2^=0.831, P<0.001). Reflecting similar patterns of sequence context described above, the trinucleotide sequence in which a mutation occurred also had a strong effect on mutability. The most mutable trinucleotides were CT<u>C</u> and CA<u>C</u>, where the final C was the mutant position (odds ratio = 3.54 and 2.02 respectively), and the least mutable were GT<u>T</u> and AG<u>A</u> (odds ratio = 0.57 and 0.58 respectively). It was not possible to combine the triplets into a single predictor, but the maximum difference in mutability between triplets indicated a strong effect of sequence context on mutability. A number of other genomic properties increased mutability, such as gene density (odds ratio =1.17) and being upstream of a transcription start site (odds ratio =1.13). Interestingly, although a change of one standard deviation in transcription level had little effect on mutability (odds ratio =1.02), the most highly transcribed sites in the genome were 3.7×more mutable than untranscribed sites.

## Discussion

In total we detected 6,843 mutations, the largest set of characterized spontaneous mutations to date. The overall rate of mutation across all lines was μ =11.5×10^−10^/site/generation, and the mutation rate for SNMs was 9.63×10^−10^ and 1.90×10^−10^ for small indels. There are therefore five SNMs for each small indel, consistent with previous results in *C. reinhardtii,* and similar to *Arabidopsis thaliana* (~5:1), but substantially lower than the ratios recently reported from MA studies in S. *cerevisiae* (33:1) and *D. melanogaster* (12:1). This large set of mutations, and the inclusion of multiple natural genotypes, allowed detailed examination of mutation rate variation between individuals within a species and mutation rate heterogeneity across the genome. In what follows we discuss the key results as they relate to the extent of mutation rate variation between natural strains and across the genome.

### Within species mutation rate variation

Our estimate of total mutation rate in *C. reinhardtii* is 14.2-fold and 4.6-fold higher than two previous estimates (Ness et al. 2012; Sung et al. 2012). The current estimate of mutation rate was partly driven by the higher rate in MA lines derived from ancestor CC-1373, but even after excluding this line the mutation rate is still substantially higher than previous estimates. The two MA lines (CC-2937-MA1, CC-2937-MA2) that were used to estimate mutation rate by Ness et al. (2012) continued to accumulate mutations for an average of ~611 generations additional generations, and the final mutation rate estimate for each of these two lines is within the confidence interval of the earlier estimate. Unfortunately, our experiment did not include strain CC-124 used in Sung et al. (2012), and so we can not directly compare mutation rates to this study. Only a single MA line (CC-1952-MA4) hada mutation rate as low as the Sung et al. (2012) estimate and the mean of all MA lines derived from that ancestor was 9 times higher. Whether this variation is the result of methodological differences or biological variation between strain CC-124 and the six strains included in our study remains to be determined.

We observed a large degree of within-species variation for the mutation rate (Fig. 1). MA lines derived from strain CC-1373 had an average mutation rate more than three times that of the other strains. MA experiments in diploid species generally start with inbred lines, and it has been argued that mutation could be affected by recessive mutation rate modifiers that are not expressed in nature. However, *C. reinhardtii* is haploid, so the elevated rate in CC-1373 must be caused by a mutation modifier that arose since collection or by natural variation expressed in nature. CC-1373 is the slowest growing of the ancestral strains, indicating that it is not well adapted to laboratory conditions. A MA experiment in *Drosophila* provided evidence that individuals in poor condition have a higher mutation rate (Sharp and Agrawal 2012), so it is possible that the higher mutation rate in CC-1373 reflects its poor condition. At the other end of the spectrum, CC-1952 had the lowest mutation rate, nearly seven-fold lower than that of CC-1373. The extent of intraspecific mutation rate variation we found implies that measuring the mutation rate for a species from a single genotype may not adequately reflect the species as a whole, and interspecific differences in mutation rate may actually reflect poor sampling within species.

In general, theory predicts that selection are expected to drive the mutation rate towards zero, because alleles that increase the mutation rate will generate deleterious alleles and thereby reduce fitness (reviewed by Sniegowski and Raynes 2013). However, mutation rates are always above zero in nature, which is usually explained by the cost of increased fidelity or by the ‘selection-drift barrier’ imposed when selection for increasingly small improvements becomes too weak to counteract genetic drift. Under both hypotheses, the extent of intraspecific mutation rate variation may reflect mutation-selection balance in genes that affect DNA-repair, replication fidelity or the susceptibility to DNA damage. In our experiment, we detected at least two MA lines with mutation rates significantly higher than their strain means ( i.e., CC-2344-MA1 and CC-2931-MA5 had mutation rates 8.0× and 3.5× above their respective strain means, Fig. 1). It is likely that these two lines acquired mutations that damaged DNA repair or stability, concordant with the presence of two mutations in DNA repair proteins in CC-2344-MA1 and five such mutations in CC-2931-MA5. However, 26 of 85 MA lines also acquired one or more mutations that affect DNA repair associated proteins, but did not have elevated mutation rates. It is possible that many of these mutations did not significantly alter the mutation rate, or that the mutations arose too late in the experiment to cause a detectable elevation of mutation rate. The increase in mutation rate in line CC-2344-MA1 was greater than the extent of natural variation among ancestral strains, suggesting that mutations that strongly alter mutation rate are common, and may segregate in natural populations until purged by selection. Therefore the high mutation rate of CC-1373 may be caused by a naturally occurring mutator allele. Alternatively, if *C. reinhardtii* is primarily asexual in nature, theory predicts that if a mutator allele results in a linked beneficial allele, the mutator will hitchhike to high frequency. A key parameter determining whether selection will favor higher mutation rates is the rate of recombination, but the frequency of sex and recombination in natural populations of *C. reinhardtii* is unknown.

### Spatial heterogeneity in mutation rate

By examining the spectrum of mutations and the local sequence in which they occur, we found clear evidence for heterogeneity in mutation rate at fine-scales. In particular, the rate of mutation at C:G sites (12.2×10’^10^) was 2.4x higher than at A:T sites (5.19×10’^10^) and transitions from C:G→T:A occurred at twice the rate expected if all mutations occurred at even rates (Fig. 2). An AT-biased mutation spectrum is consistent with a growing body of evidence suggesting that it might be universal in prokaryotes (Hershberg and Petrov 2010) and eukaryotes (e.g. Zhu et al. 2014). Additionally, we found that the sequence flanking a mutated site strongly influenced the mutation rate. In mammals methylated CpG sites are frequently deaminated, causing C to T transitions, but in *C. reinhardtii* there is only weak evidence of CpG methylation, and our data reveals only a small excess of CpG motifs in C to T mutations (Fig. 3). The most mutable triplet (CTC) had a mutation rate more than 10× higher than the least mutable triplet (GCA), and after accounting for background triplet frequencies, a mutation from CTC to CTT was 17× more likely than a mutation from AAA to AAG. Interestingly, this CTC triplet appears to be highly mutable across a very wide diversity of organisms, including fungi (Zhu et al. 2014), plants and animals (Alexandrov et al. 2013b). In human tumor genomes, there is a predominance of C to T and C to G mutations in the same CTCG sequence motif, which has been linked with the APOBEC family of cytidine deaminases (Alexandrov et al. 2013b). Given that this motif has been found repeatedly, it seems probable that the mutability of other sequence motifs may be shared across species, however the mechanisms underlying this phenomenon are unknown. The fact that the mutation rate can vary to this extent over very short scales has consequences for the evolution of DNA and protein sequence. In the future, incorporation of direct measurements of mutability into models of sequence change will facilitate better predictions of disease susceptibility and molecular evolution (see Michaelson et al. 2012; Neale et al. 2012; Samocha et al. 2014).

By comparing the distribution of intermutation distances to a random expectation, we found that there is an excess of mutations clustered within 1-10bp of one another (Fig. 4). The fact that these clusters all occur within MA-lines suggests that each represents a single multinucleotide mutation (MNM) event. In total there were 80 pairs and two trios of MNMs within 10bp of one another, implying that 2.8% of SNMs arise through clustered mutations. The average proportion of MNMs was similar in MA studies of S. *cerevisiae, D. melanogaster, C. elegans* and *A. thaliana* (3.4%), and genome sequencing of humans (1-4% Schrider et al. 2011; Harris and Nielsen 2014). The generation of these clusters has been linked to error prone polymerases such as Pol ζ in S. *cerevisiae* (Stone et al. 2012; Northam et al. 2013). In human and S. *cerevisiae* the Pol ζ enzyme creates an excess of GC to AA or TT MNMs (Northam et al. 2013; Harris and Nielsen 2014). Although we did not observe a similar excess of mutations at GC sites, we found that 27 of 55 dinucleotide MNMs occur at CC sites and that 25 of these resulted in AA/AT/TA/TT dinucleotides. The consistency of these results across taxa suggests that we cannot consider MNMs as an oddity of the mutational process. MNMs violate the assumption of independence between SNP sites and could potentially lead to mis-inferences about the nature of selection in the genome. Additionally, by altering two or more nearby sites, MNMs have the potential to move between fitness peaks that would otherwise require maladaptive single mutations as intermediates.

At large genomic scales, we found little evidence for heterogeneity of the mutation rate. For example, the mutation rate variation among 200Kbp windows could be largely accounted for by random fluctuations. Although we found clear evidence of fine-scale variation in mutation rate, the variation appears to be evenly spread along the chromosome. This effect can be seen in our predictive model of mutation, where the mutability of sites in 200Kbp windows averages out, so that the standard deviation among windows equates to ~7.5% of the mean (i.e., mean mutability = 0.069, SD = 0.005). Our findings are consistent with direct measurements of mutation rate in *D. melanogaster* (Schrider et al. 2013), S. *cerevisiae* (Zhu et al. 2014) and humans (Kong et al. 2012), where no evidence of large scale variation in the mutation rate was detected. Although, comparative evidence suggests that substitution rate varies at the scale of megabases in mammals, this may be driven by selection or biased gene conversion during recombination. From our observations and direct estimates of mutation rate variation in other species, we conclude that the causes of mutational heterogeneity do not appear to operate at the scale of kilobases, and if heterogeneity exists at this scale it will require even more precise measurements of the mutation rate.

### Factors that predict mutability

Our model of mutability identified a number of other genomic properties that predict the rate of spontaneous mutation and create heterogeneity between sites. For example, the %GC of the 10bp around a mutated site was positively correlated with mutability (Odds ratio = 1.38, 1-SD=16.3%), probably because G:C bases and GC-rich triplets were more mutable. However, the GC-content of the 1,000bp surrounding a site was negatively associated with its mutability (e.g., %GC of 1,000bp window, Odds ratio = 0.12, 1-SD=5.4%). A negative correlation between mutability and GC content in humans has been attributed to higher melting temperatures of GC-rich DNA (Fryxell and Moon 2005). Because cytosine deamination is one of the most common sources of mutation and only occurs while DNA is single stranded, mutation is less common in regions with high melting temperature (Frederico et al. 1993). An alternate explanation for our observations is that sites with a high mutation rate, for an unknown reason, evolve low GC-content because mutation is AT-biased.

Our model of mutability also revealed an effect of gene expression when comparing untranscribed DNA to the most highly transcribed genes (odds ratio = 3.71). However, because most regions are untranscribed and the variance of transcription in expressed genes is relatively low, transcription level overall had little effect on mutability (odds ratio 1.02, 1-SD=108.3 FPKM). It is commonly reported that highly expressed genes are the most evolutionarily conserved, therefore an elevated mutation rate would predict that more deleterious mutations should occur in high expression genes and therefore more purifying would be required to conserve these sequences. The mean mutability score varied across sites with different annotations. The 5’ and 3’ UTRs had the highest mutability (predicted μ = 1.5x10^−9^), which is consistent with the observation in other species that these regulatory regions are often found in open chromatin, allowing binding of transcription factors, and potentially leading to more damage to the DNA. Consistent with an increased mutation rate and AT-biased mutation, UTRs have the lowest GC content of any broad category of sites (56.7%). Although the model predicted a higher rate in UTRs, we did not observe an elevation in observed mutation rate, possibly because even with nearly 7,000 mutations there was still insufficient power to detect such subtle variation. Overall, the model accurately predicted the observed mutation rate, demonstrating that average mutation rate can be predicted from key genomic properties (Fig. 5). However, variation in mutability may not be fully captured with this approach Eyre-Walker and Eyre-Walker (2014). For a close fit between observed and predicted mutability, only the average mutability of each bin needs to be accurately predicted. There may still be unexplained variation around the mean within each bin and we should be cautious about predictions of mutability for very small numbers of sites. However, for large groups of sites the model accurately predicts the average mutation rate and we can be confident in the genomic properties that best predict mutation rate. Mutability also revealed that mutation rate variation affects patterns of neutral genetic variation. We found a clear positive relationship between mutability and nucleotide diversity at silent sites (Fig. 6). The model identifies the genomic properties of sites that mutated in our experiment and we show that using these genomic properties we are able to predict natural levels of genetic diversity. This implies that our model captures the variation in mutation rate that exists under natural conditions.

This study characterized the largest set of spontaneous mutations to date and demonstrated the insights that can be gained by combining MA with whole genome sequencing. We found 7-fold variation in mutation rate among natural strains of *C. reinhardtii.* Although the mutation rate did not vary across large genomic windows, the mutation rate of individual sites was strongly affected by their flanking sequence, resulting in fine-scale heterogeneity of mutation rate. Other genomic properties, such as GC content, gene density and expression level, also influenced mutability. Similar results across a wide diversity of species suggests that general properties of mutation exist and that models of sequence evolution could be improved to reflect these properties and better detect selection in the genome or estimate phylogenetic relationships. In the near future rapidly evolving sequencing technologies will facilitate even more detailed investigation into the process of mutation from both MA and parent-offspring sequencing. One important avenue of future research will be a synthesis of findings from studies like ours with the underlying DNA repair and damage mechanisms to provide explanations for patterns mutational heterogeneity between individuals and across the genome.

## Methods

### Mutation accumulation experiment

We conducted a mutation accumulation experiment in six genetically diverse wild strains of *C. reinhardtii* obtained from the Chlamydomonas Resource Center (chlamycollection.org). The strains were chosen to broadly cover the geographic range of known *C. reinhardtii* samples in North America (Table 1). To initiate the MA lines, a single colony from each of the six ancestral strains was streaked out, and we randomly selected 15 individual colonies to start the replicated MA lines (for a total of 90 MA lines). We bottlenecked the MA lines at regular intervals by selecting a random colony which was streaked onto a fresh agar plate. We estimated the number of generations undergone by each MA line over the course of the experiment by measuring the number of cells in colonies grown on agar plates after a period of growth equivalent to the times between transfers in the experiment. The details of the MA line creation and generation time estimation can be found in Morgan et al. (2014).

### Sequencing and alignment

To extract DNA, we grew cells on 1.5% Bold’s agar for 4 days until there was a high density of cells, at which point the cells were collected and frozen at −80°C. We disrupted the frozen cells using glass beads, and extracted DNA using a standard phenol-chloroform extraction. Whole-genome re-sequencing was conducted using the Illumina GAII platform at the Beijing Genomics Institute (BGI-HongKong Co., Ltd, Hong Kong). The sequencing protocol was modified to accommodate the unusually high GC content of the *C. reinhardtii* genome (mean GC= 63.9%). Variation in GC-content is known to cause uneven representation of sequenced fragments, especially when GC > 55% (Aird et al. 2011). We therefore used a modified PCR step in sequencing library preparation, following Aird et al. (2011) (3 min at 98°C; 10 × [80 sec at 98°C, 30 sec at 65°C, 30 sec at 72°C]; 10 min at 72°C, with 2M betaine and slow temperature ramping 2.2°C/sec). We obtained ~30× coverage of the genome (3Gbp of 100bp paired-end sequence) for each of the MA lines.

We aligned reads to the *C. reinhardtii* reference genome (version 5.3) using BWA 0.7.4-r385 (Li and Durbin 2009). We included the plastid genome (NCBI accession NC_005353), mitochondrial genome (NCBI accession NC_001638) and the MT-locus (NCBI accession GU814015) to avoid misalignment of reads derived from these loci onto other parts of the nuclear genome. We tested a variety of values for the fraction of mismatching bases allowed in alignments, but variation about the default (n=0.04) did not improve the number of high quality reads mapped or genome coverage (results not shown). After alignment, we removed duplicate reads with the Picard tool MarkDuplicates (v1.90). To avoid calling false variants due to alignment errors, we used the GATK (v2.8-1) tools RealignerTargetCreator and IndelRealigner (Mckenna et al. 2010; Depristo et al. 2011) to realign reads flanking potential insertions and deletions. We realigned all replicate MA lines from each starting strain together to ensure that the same alignment solutions were chosen in all lines derived from that strain. The realigned BAM files included all MA lines from given ancestral strain and were then used to jointly call genotypes using the UnifiedGenotyper from GATK. We used the “--output_mode EMIT_ALL_SITES” option to output all genomic positions so that we could identify both high quality sites regardless of whether they had mutated. We used a “heterozygosity” parameter of 0.01, but previous testing in *C. reinhardtii* showed that our genotyping is not sensitive to this prior as long as read depth is high, as it is in the present experiment (Ness et al. 2012). To identify short insertions and deletions (indels) we used the GATK v(2.8-1) tool ‘HaplotypeCaller’, which performs local re-assembly of reads (i.e., indels called with UnifiedGenotyper were ignored). The six resulting Variant Call Format files (VCFs) (one per ancestral strain) were converted to wormtable databases using the python package WormTable v0.1.0 (Kelleher et al. 2013) which enabled efficient exploration of quality filters for mutation identification.

### Mutation identification

MA lines within an ancestral strain were genetically identical at the start of the experiment, so any unique allele carried by a replicate within a strain was a candidate mutation. We applied a number of filters to genotype calls to identify mutations, while minimizing false positive and false negative calls. A site was called as a mutation if within that ancestral strain:

1. The mapping quality (MQ) ≥ 90 and the PHRED called site quality (QUAL) ≥ 100
2. All MA lines were ‘homozygous’; *C. reinhardtii* is haploid therefore this filter avoided mapping errors due to paralogous loci.
3. The genotype of exactly one MA line differed from the rest of the lines
4. All non-mutated lines shared the same genotype
5. At least two sequences have confident genotype calls

### Callable sites

To calculate mutation rates and define null expectations, we needed to know the total number of sites with equivalent quality to the new mutations, hereafter referred to as "callable” sites. However, the definitions and distributions of quality scores are often different for variant and invariant sites. We therefore inferred a second measures of quality for invariant sites that was comparable to that used for mutant sites. For each mutant site we extracted the QUAL and MQ for the mutation and the nearest invariant site, under the assumption that because most reads are shared between adjacent sites the quality characteristics of the sites will be similar. We then estimated the correlation and relationship between quality scores at neighboring mutant and invariant sites using a linear model (MQ: R^2^=0.9996, *P<* 0.001, QUAL: R^2^=0.38, *P<* 0.001). The linear relationships between invariant and variant quality scores were used to predict appropriate MQ and QUAL thresholds for invariant sites (invariant MQ threshold = 90, invariant QUAL threshold =36.4). Analogous to the mutation calling, a site was callable within an ancestral strain if no line was called as a heterozygote, all lines with mapped reads had the same genotype call and at least two MA lines had genotype calls.

### Sanger confirmation

We estimated the accuracy of our mutation calls using Sanger sequencing. We randomly selected 192 mutation calls (32 per ancestral strain) including both short indels and SNMs. We amplified each locus in the putative mutant MA line and a non-mutated MA line from the same ancestral strain. Sequences were then visually inspected in SeqTrace v0.9.0 to confirm the presence of the mutated site.

### Mutation rate calculations

We calculated the mutation rate (μ) in each replicate as, μ = mutations / (callable sites χ MA generations). Whenever multiple MA lines were combined for mutation rate calculations, the number of callable sites and MA generations (site-generations) for each MA line was included to accurately account for differences amongst replicate lines. Similarly, all null expectations and mutation rate estimates for particular classes of sites take into account the number of site-generations for the specific positions included. To compare the average mutation rate of the six ancestral strains, we used the GLS function in R to fit a linear model to the individual mutation rate estimates of the MA lines. The model included mutation rate as the response variable and ancestral strain as a fixed factor. We allowed the variance to differ among ancestral lines using the varIdent function (Zuur et al. 2009). We then used the ghlt function to generate linear contrasts, allowing us to further explore differences among the ancestors.

### Base composition and sequence context

Throughout our analyses of the mutation spectrum, we treated complementary mutations (C:G and A:T) symmetrically, such that there were six distinct SNMs (A:T→C:G, A:T→G:C, A:T→T:A, C:G→A:T, C:G→G:C, C:G→T:A). To assess the base spectrum of mutations, we calculated the frequency of each of the six mutation types relative to the expected frequency calculated from the base composition of the callable sites. To analyze the local sequence context in which mutations occurred, we measured base composition at each of the positions 5bp upstream and downstream of the mutated site. To calculate the null expectation for sequence context we estimated base composition in analogous windows surrounding 10^6^ randomly selected callable sites. Separate expectations were generated for sites centered on A:T and C:G.

### Spatial heterogeneity of mutation

To assess whether there was spatial heterogeneity in mutation rate we calculated the mutation rate across the genome in sliding windows. We conducted the analysis with windows of 100Kbp, 200Kbp, 500Kbp and 1Mbp but because the results were qualitatively similar and we report only the 200Kbp analysis. The mutation rate of each window was calculated as the number of mutations in that window divided by the total number of callable site*generations. To assess how the mutation rate in these windows varied relative to null expectations, we simulated a random distribution of mutations. For each MA line we generated a corresponding simulated line where the number of mutations carried by that line was distributed amongst the 200Kbp windows in proportion to the number of callable site-generations in each window. This procedure was repeated 1,000 times to generate an expected distribution of mutation rates across the 200Kbp windows.

We also tested for the presence of a non-random spatial distribution of mutations by comparing the observed distribution of intermutation distances to a simulated distribution. This approach differs from the analysis above because it can detect fine scale clusters of mutations. We simulated data under a model where mutations occur randomly across the genome, while retaining the same number of mutations per MA line and accounting for differences in the callable genome positions. For each MA line we generated a corresponding simulated sample by randomly assigning the number of mutations that occurred in that MA line to individual callable positions. This allowed us to assess whether there was significantly more clustering within and between lines while accounting for line-specific differences in callable sites. The observed and simulated distributions of intermutation distances were compared using the Kolmogorov-Smirnov (KS) test in R.

### Mutability

To determine which genomic properties influenced the mutability of individual sites we used regularized logistic regression to differentiate between the identified mutations and randomly selected callable sites. Our analysis was loosely based on the approach of Michaelson et al. (2012). For all 6,843 mutations and 10^5^ non-mutated sites, we collated a table of genomic properties and annotations to use as predictors in the logistic regression. Genomic properties included %GC, gene density, transcription level, recombination rate, nucleosome occupancy and the trinucleotide sequence in which the site occurs (see Supplementary table S1 for details). A number of genomic properties were calculated for each site in windows of varying size from 10bp up to 1Mbp. Categorical predictors were converted to multiple binary predictors (0/1 for each category level) to be fitted in the same model with numeric predictors.

With these predictors we used the R package GLMnet (v1.9-8) (Friedman et al. 2010) to fit a logistic regression, where mutation class (mutant (1) or background (0)) was the binary response variable. From the model, we estimated mutability at each site in the genome as its probability of belonging to class ‘mutation’ given the genomic predictors at a given site. We assessed the accuracy of the predicted mutability by binning sites into 100 uniformly spaced mutability categories from 0.0-1.0. The exact value of the mutability score was not relevant, since it only reflected the ratio of mutations to background sites in the original training set. Within each mutability category the number of observed mutations divided by the total number of site-generations in that category was used to estimate the mutation rate. The observed mutation rate was predicted to be positively correlated with the mid-point mutability of the category. To test whether mutability predicted long term effects of mutation rate variation, we also calculated the relationship between mutability and natural levels of nucleotide diversity in the six ancestral strains used to start the MA lines. In neutrally evolving haploid DNA the level of nucleotide diversity (θ_π_) is expected to be twice the product of mutation rate and the effective population size (*2N_e_μ),* we therefore predict that the mutation rate should correlate positively with mutability. For this analysis whether a site was variant was omitted from the model in order to avoid circularity in the relationship between diversity and mutability. We binned silent sites (intergenic, intronic and 4-fold degenerate sites) into 100 uniformly spaced mutability categories from 0.0-1.0 and calculated θ_π_ for all sites in each bin.

To assess the relative contributions of each genomic property to mutability, we extracted the coefficients of each predictor from the model. GLMnet can handle highly correlated predictors using an elastic-net penalty *(α)* that will either shrink coefficients toward each other (ridge penalty, α=0) or keeps one and discard the others (lasso, *α*=1). The fit of the model was unchanged by the selection of α and all results presented used *α*=0.01. To compare the log(odds ratio) of each genomic property on mutability, we scaled each predictor so that a change from 0.0 to 1.0 was a change of one standard deviation. As alternate scaling we also normalized the predictors such that each ranged from exactly zero to one.

### Data Access

All genomic data generated as part of this project is publicly available through the NCBI Sequence Read Archive (SRA BioProject Accession: SRP052900)

---

**Supplementary Figure S1.**
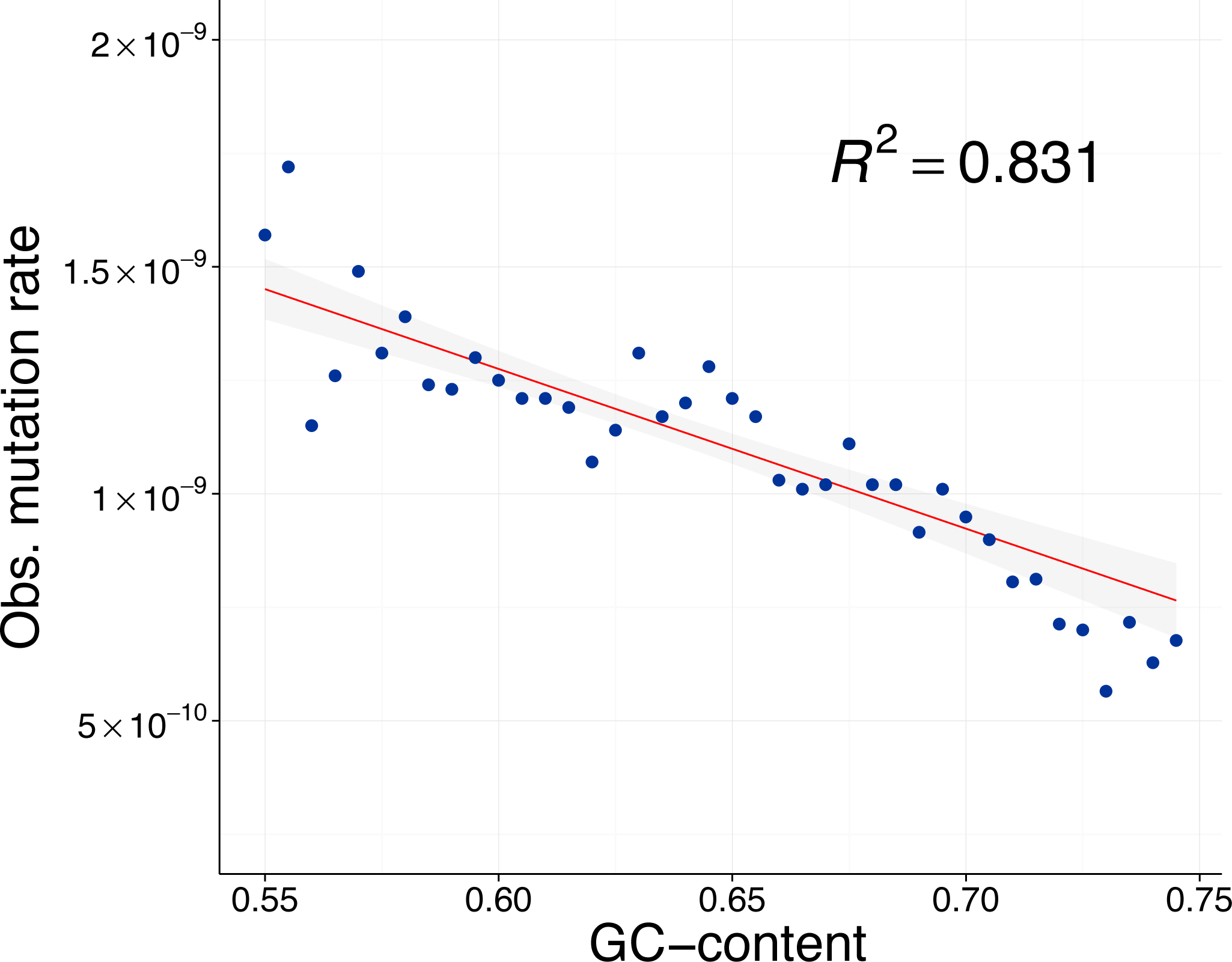
Relationship between mutation rate and GC content. Linear fit between observed mutation rate and GC content of the 1000bp surrounding a site. Each point represents multiple genomic sites placed in discrete bins (width = 0.005) based on each site’s GC content. Observed mutation rate for each point was calculated as the number of observed mutations divided by the total number of callable sites-generations in that bin. The shaded grey area around the line represents the 95% confidence region.

